# Natural infection of SARS-CoV-2 delta variant in Asiatic lions (*Panthera leo persica*) in India

**DOI:** 10.1101/2021.07.02.450663

**Authors:** Anamika Mishra, Naveen Kumar, Sandeep Bhatia, Ashutosh Aasdev, Sridhar Kanniappan, A. Thayasekhar, Aparna Gopinadhan, R. Silambarasan, C. Sreekumar, Chandan Kumar Dubey, Meghna Tripathi, Ashwin Ashok Raut, Vijendra Pal Singh

## Abstract

In May 2021, severe acute respiratory syndrome coronavirus 2 (SARS-CoV-2) was detected in nine Asiatic lions (*Panthera leo persica*) in Arignar Anna Zoological Park, Chennai, Tamil Nadu, India. Sequence and phylogenetic analysis showed that the SARS-CoV-2 viruses belong to a variant of concern (VOC, delta variant, B.1.617.2 lineage) and that these viruses clustered with B.1.617.2 lineage viruses of the same geographical region detected in the same month.

---

Since the first identification of severe acute respiratory syndrome coronavirus 2 (SARS-CoV-2) in 2019 in the city of Wuhan, Hubei province, China, its associated disease coronavirus disease 2019 (COVID-19) has resulted in 176,531,710 confirmed human cases, with 3,826,181 deaths as of June 17, 2021 (Zhou et al., 2020; World Health Organization, 2021). Because viruses closely related to SARS-CoV-2 have been found in bats and pangolins, they are being debated as possible SARS-CoV-2 origins (Lam et al., 2020; Liu et al., 2020; Zhang et al., 2020). For the reason that SARS-CoV-2 is believed to have a zoonotic origin, finding susceptible animal species, reservoirs, and cross-species transmission events is a subject of global scientific and public interest. Experimental studies showed that SARS-CoV-2 infects and replicates efficiently in domestic cats, ferrets, and fruit bats but poorly in dogs. However, chickens, ducks, and pigs do not seem to support productive SARS-CoV-2 infection (Shi et al., 2020; Kim et al., 2020). Conversely, in natural conditions, SARS-CoV-2 seems to have broader host susceptibility range and as of 31 May, 2021, SARS-CoV-2 infections has now been reported in at least ten animal species, namely, cats, dogs, tigers, lions, snow leopards, pumas, gorillas, pet ferrets, otter and farmed mink from thirty countries resulting in 552 outbreaks worldwide (OIE, 2021). Importantly, in most of these natural infection cases, caretakers or owners of these animal species have been found positive for SARS-CoV-2 infection, pointing the transmission of infection from humans to animals. In rare cases, two way transmissions (transmission of infection from humans to animals, and back to humans) have also been documented in farmed minks in the Netherlands (Oude Munnink et al., 2021).

In this study, we report the first confirmed natural SARS-CoV-2 infections in Asiatic lions (*Panthera leo persica*) caused by a delta variant as per World Health Organization (WHO) nomenclature (PANGO lineage B.1.617.2, Nextstrain clade 21A) in India, and provide a thorough genomic characterization of SARS-CoV-2 obtained from the infected Asiatic lions. These confirmed natural infections in the India occurred in last week of May, 2021, when India was battling an unprecedented surge in COVID-19 cases.

The International Union for Conservation of Nature (IUCN) has classified Asiatic lions as Endangered, and their population is steadily declining owing to poaching and habitat degradation (Bauer et al., 2016). Some Asiatic lions in Arignar Anna Zoological Park, Chennai, Tamil Nadu, India experienced loss of appetite, nasal discharge and occasional coughing symptoms in the last week of May 2021. COVID-19 claimed the lives of two lions, Neela and Pathbanathan, on June 3 and 16, 2021, respectively. Nasal swabs, rectal swabs, and faecal samples were taken from eleven Asiatic lions (6 males and 5 females) during 24 to 29 May, 2021 and sent to the ICAR-National Institute of High Security Animal Diseases in Bhopal, India, for molecular investigations (Table S1). The genome of SARS-CoV-2 was confirmed in nine lions using the VIRALDTECT - II Multiplex Real Time PCR Kit (GENES2ME Pvt. Ltd., India), which enabled identification of SARS-CoV-2 E, N, and RdRp genes. Furthermore, all of the lion samples tested negative for Canine Distemper Virus using the OIE-recommended RT-PCR method (Frisk et al., 1999).

Subsequent to confirmation of SARS-CoV-2 infection in the lions, whole genome sequencing of SARS-CoV-2 was generated directly from nasal swabs from four lions (Lioness-Jaya, Lion-Shankar, Lioness-Niranjana, and Lion-Pradeep). In a nutshell, tiling PCR spanning over the whole genome of SARS-CoV-2 was carried out with the Artic network primers (https://artic.network/). Cleaning and quantification of PCR products were performed and 100ng of each sample was used to create a barcoded sequencing library using the PCR Barcoding Kit (SQK-PBK004). The sequencing was done on an Oxford Nanopore Technologies MinION with a R9.4.1 flowcell, yielding a total of 300MB of data. To assemble the whole genome, guppy v5.0.7 (Oxford Nanopore Technologies) was used for base calling and de-multiplexing, followed by Porechop (https://github.com/rrwick/Porechop) for adaptor removal. The readings were mapped to the reference SARS-CoV-2 genome (NC 045512) using minimap2 v2.17 (r941) (Li, 2018), and the variations were called using nanopolish v0.13.2 (Loman et al., 2015). After two rounds of nanopolish, four complete genomes were produced and submitted to NCBI with the GenBank accession numbers MZ363851-MZ363854.

To better understand the temporal dynamics of SARS-CoV-2, complete SARS-CoV-2 genomes with high coverage originating from the Tamil Nadu state of India (n = 310) during 1 January 2021 to 11 June 2021 were downloaded from GISAID on 11 June 2021 (Elbe and Buckland-Merrett, 2017, https://www.gisaid.org/). These genomes were clustered at 99.9% identity threshold using UCLUST algorithm to generate a set of representative sequences (Edgar, 2010). PANGO lineages (Rambaut et al., 2020) and Nextstrain clades (https://clades.nextstrain.org/) were assigned to these representative sequences. Along with the SARS-CoV-2 genomes in lions of this study and Wuhan-Hu-1 reference sequence (EPI_ISL_402124), these representative sequences were aligned using MAFFT v.7.475 (Katoh and Standley, 2013) and a phylogenetic tree was constructed using the GTR-Gamma model in RAxML v. 8.2.12 (Stamatakis, 2014). In addition, complete SARS-CoV-2 genome having high coverage from the lions (n = 24) across the globe were downloaded from GISAID on 11 June 2021 and a phylogenetic tree was constructed using the GTR-Gamma model in RAxML v. 8.2.12 (Stamatakis, 2014).

Compared to the Wuhan-Hu-1 reference sequence (EPI_ISL_402124), all the four Asiatic lions sequences contained 24 amino acid substitutions and 2 deletions. The tempo-spatial dynamics of substitutions across the multiple proteins encoded by the SARS-CoV-2 detected in lions, as well as their functional roles are provided in Table S2. The spike protein sequence of SARS-CoV-2 in lions had 9 amino acid substitutions and 2 deletions compared with the Wuhan-Hu-1 strain, and are typically matched with a delta variant (PANGO lineage B.1.167.2) of SARS-CoV-2 viz., N-terminal domain (NTD; T19R, G142D, E156del, F157del, R158G), receptor binding motif (RBM; L452R, T478K), Subdomain 2 (SD2, D614G), substitution close to S1/S2 protease cleavage site (P681R), and heptad repeat 1 (HR1, D950N) (Figure 1). Additionally, they carried the K77T substitution in the NTD, which has been reported from 24 countries, including India, with a low frequency of 0.44%, but it occurs 27.42 % of the time (65/237 sequences) in the lineage in Tamil Nadu, India (Table S2). SARS-CoV-2 had previously been documented to cause natural infections in lions from the United States (B, 19A), Spain (B.1.177), and the Czech Republic (B.1.1.7, Alpha), as well as two lineages in tigers from the Czech Republic (B.1.1.7, Alpha) and the United States (B.1) (Figure S1) (McAloose et al., 2020). As a result, our study describes the first confirmed natural SARS-CoV-2 infection in Asiatic lions in India, which was caused by a variant of concern (VOC delta, B.1.167.2 lineage).

**Figure 1.**
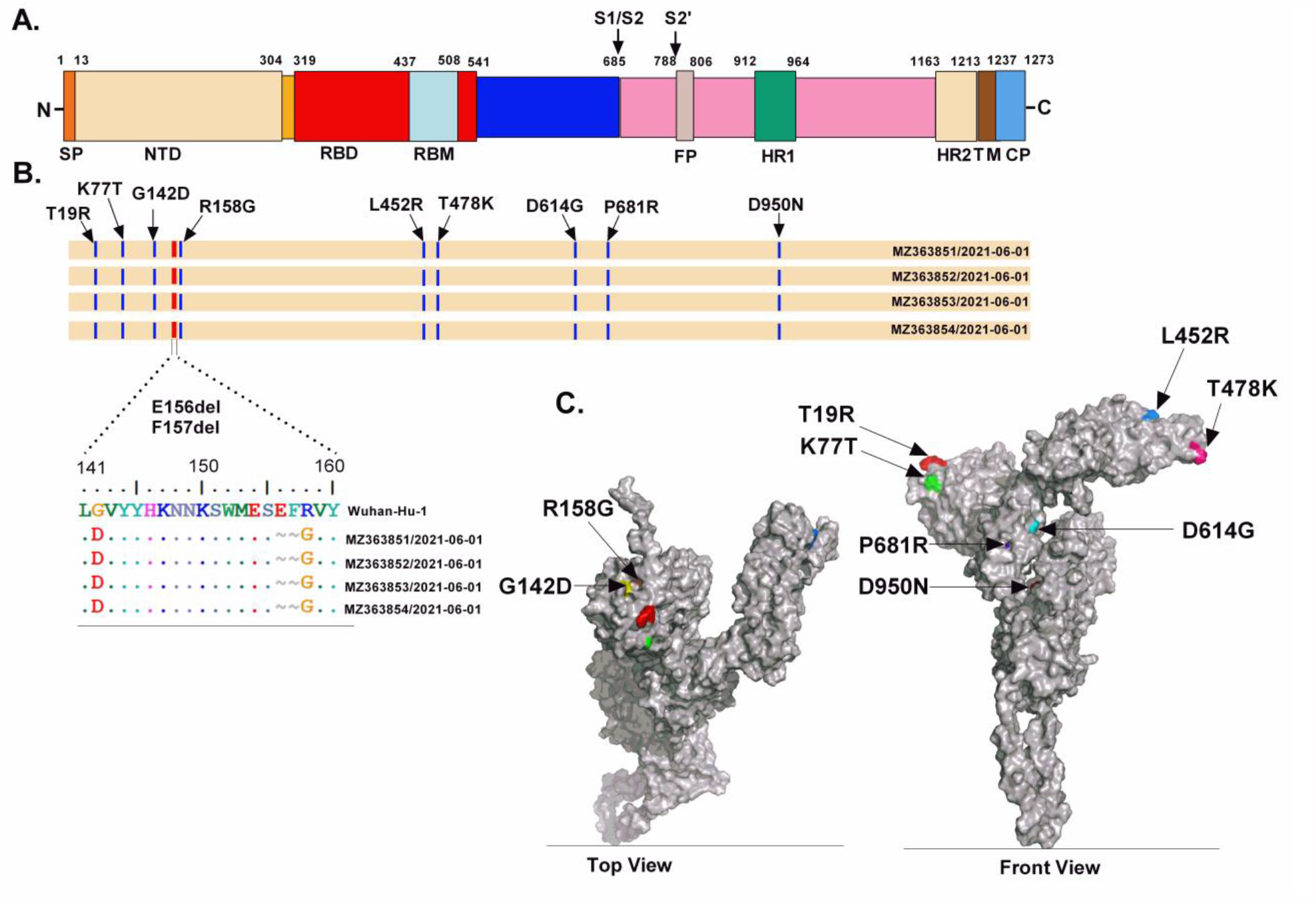
(A) shows schematic representation of functional domains of the spike (S) protein for SARS-CoV2 Wuhan-Hu-1; (B) A comparison of amino acid changes detected in the S protein of SARS-CoV-2 in Asiatic lions with respect to Wuhan-Hu-1 reference sequence (EPI_ISL_402124); (C) Mapping of amino acid substitutions noted in Asiatic lion SARS-CoV-2 (GenBank accession no. MZ363851) on to the structural model of S protein, build using I-TASSER (Iterative Threading ASSEmbly Refinement) on a template (PDB: 6acc) (Yang et al., 2015).

The spike (E156del, F157del, and R158G) and NS3 (V88I) deletions and substitutions noted in all four lion sequences were not found in human virus sequences from the same geographic location. This could be due to insufficient and small-scale sequencing of SARS-CoV-2 in this geographic region, causing the prevalence of these deletions and substitutions to be underestimated. Although NS3 (V88I) substitution frequency is very low in India as well as world, the B.1.36.8 lineage in Punjab, a northern state of India carried this substitution in July 2020 (Table S2). It is possible that NS3 (V88I) was inserted at random during virus replication. Furthermore, these deletions and substitutions in spike and NS3 were not seen in previously reported lion sequences (from the United States, Spain, and the Czech Republic), ruling out the possible host-adapted mutations. However, experimental investigations are needed to see if deletions and substitutions in spike protein, namely E156del, F157del, R158G, and K77T, are escape mutants or related with increased transmissibility and/or pathogenicity.

Blasting of all the four lions viral genome sequences against the available sequences in the GISAID closely matched with a SARS-CoV-2 sequence (GISAID accession no. EPI_ISL_2463770) found in a patient belonging to the same geographical region in May 2021. Furthermore, phylogenetic analysis of viral genome sequences from the lions, as well as all the sequences available from Tamil Nadu state of India, clustered them into the human B.1.617.2 lineage (VOC Delta, GISAID clade - G/478K.V1, Nextstrain clade - 20A/S:478K) (Figure S2). Importantly, viral genome sequences from all the lions closely matched a representative SARS-CoV-2 sequence (GISAID accession no. EPI_ISL_2463770, which represents 152 viral genome pools of the same geographical region), corroborating with the lions’ sample collection month (Figure 2). In the absence of any recent new animal introductions to the zoo, no public access to the zoo during COVID-19 imposed strict lockdown and consistent use of personal protective equipment (PPE) by the zoo keepers, the source of SARS-CoV-2 infection in the Asiatic lions cannot be ascertained and can only be presumed to be via unpremeditated contaminated fomites. Seven of nine infected lions belong to the lion safari of the zoo and hence share a common habitat including shelter, food space and water source; the other two lions are display animals who share a common moat. In the presence of shared habitat offering several opportunities of close physical contact, identification of infection with genetically identical SARS-CoV-2 in these lions in short period of time, clearly indicates the likelihood of lion-to-lion transmission, which might be of grave concern.

**Figure 2.**
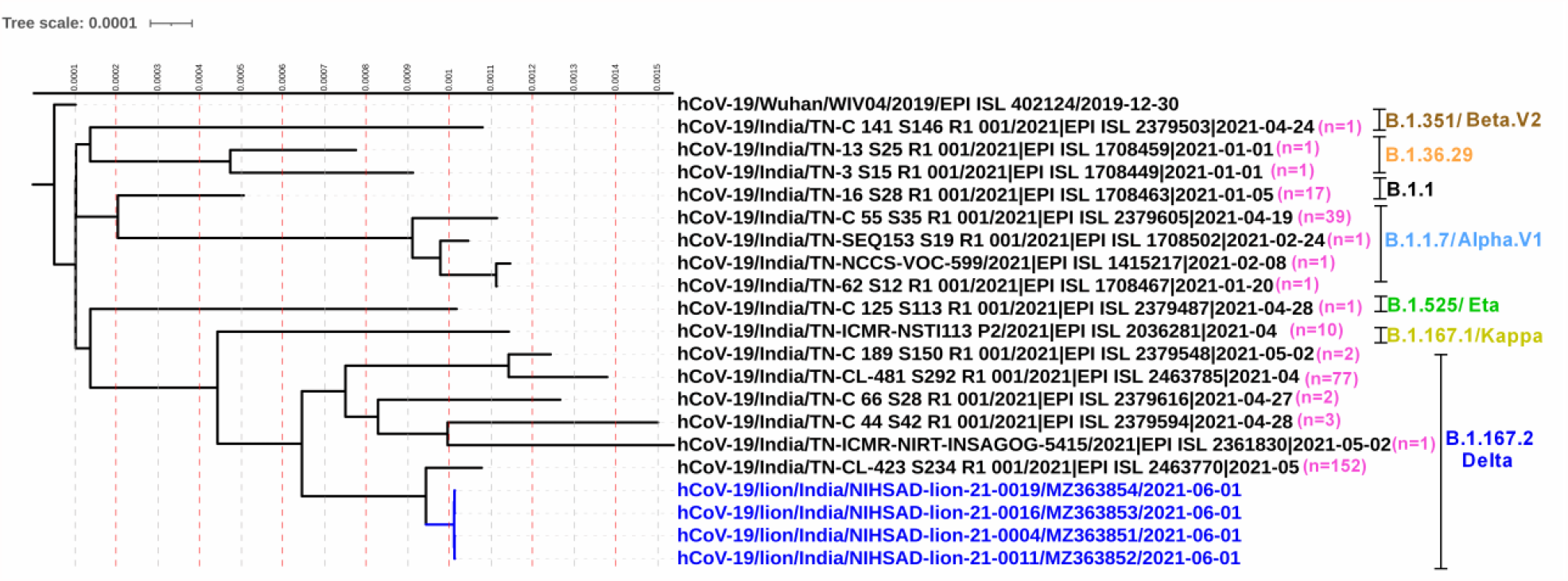
Complete genome phylogenetic analysis of Asiatic lions SARS-COV-2 sequences with Wuhan-Hu-1 reference sequence (EPI_ISL_402124) and representative sequences of different clusters (generated at 99.9% identity threshold) from the available SARS-COV-2 sequences from Tamil Nadu state of India. The Maximum Likelihood (ML) tree was rooted to Wuhan-Hu-1. The number in parentheses indicates the number of SARS-CoV-2 genome sequences clustered at 99.9% identity threshold.

In conclusion, we reported the first confirmed natural SARS-CoV-2 infections in Asiatic lions (*Panthera leo persica*) in India, which were caused by a delta variant (B.1.617.2 lineage). This finding justifies increased surveillance for this VOC in other wild species, as well as strict biosecurity measures to check sick/asymptomatic handlers/keepers/visitors from entering the area.

## Acknowledgement

We, the authors, are thankful to the Director, Arignar Anna Zoological Park, Chennai, India, for sending the Asiatic lion’s samples for SARS-CoV-2 molecular investigations and to Veterinary officer, Arignar Anna Zoological Park, Chennai, India for providing the clinical information of lions. We are grateful to the Director, ICAR-NIHSAD, Bhopal for providing infrastructural facilities and funding for this study. We appreciate Dr. Atul Kumar Pateriya’s assistance with RT-qPCR. We also acknowledge the originating and the submitting laboratories for sharing the SARS-CoV-2 genomic sequence data via the GISAID.

---

**Figure S1.**
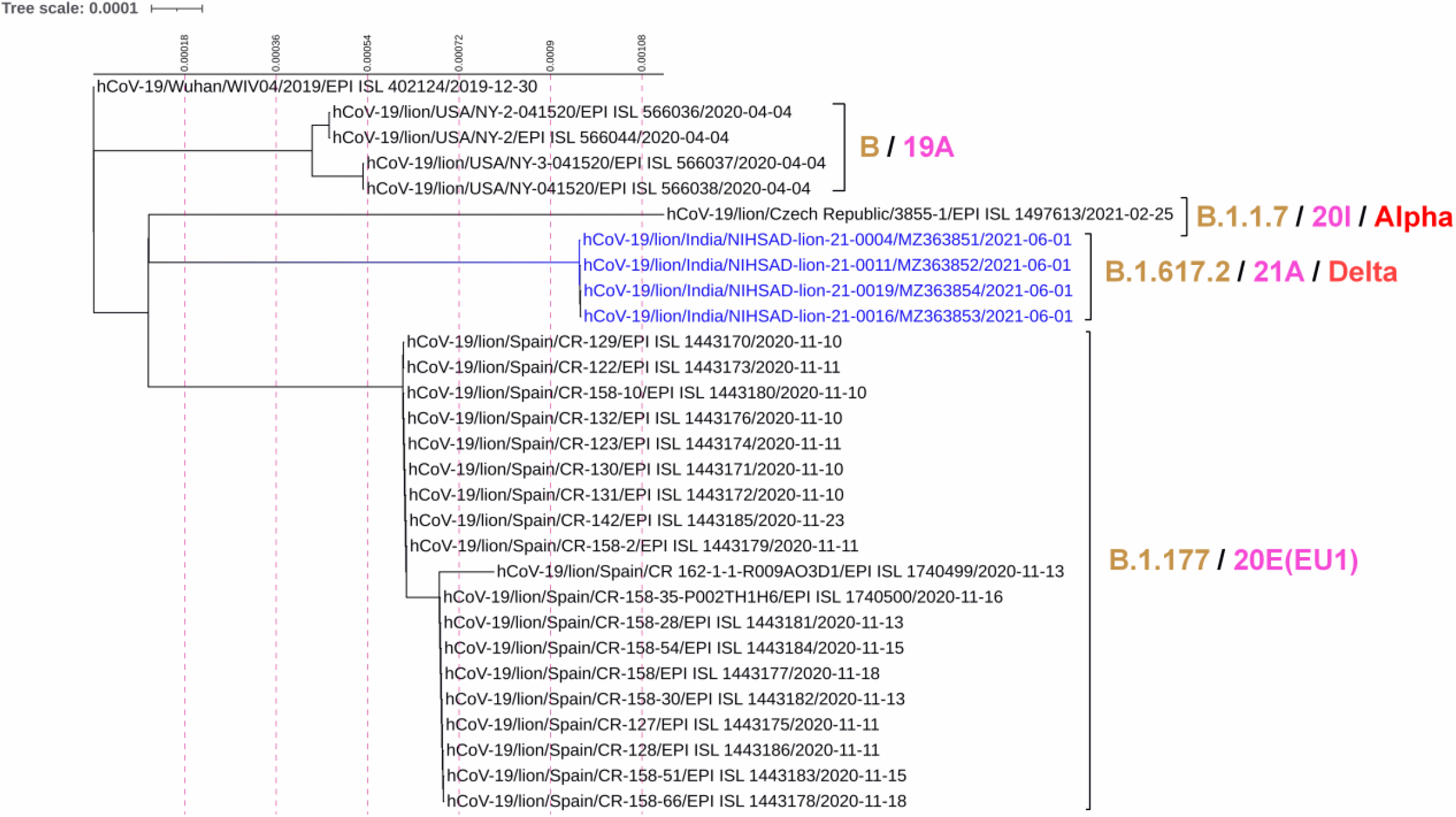
ML tree showing the phylogenetic relationship among the Asiatic lions SARS-COV-2 sequences of this study with other available lions and tigers sequences in the GISAID.

**Figure S2.**
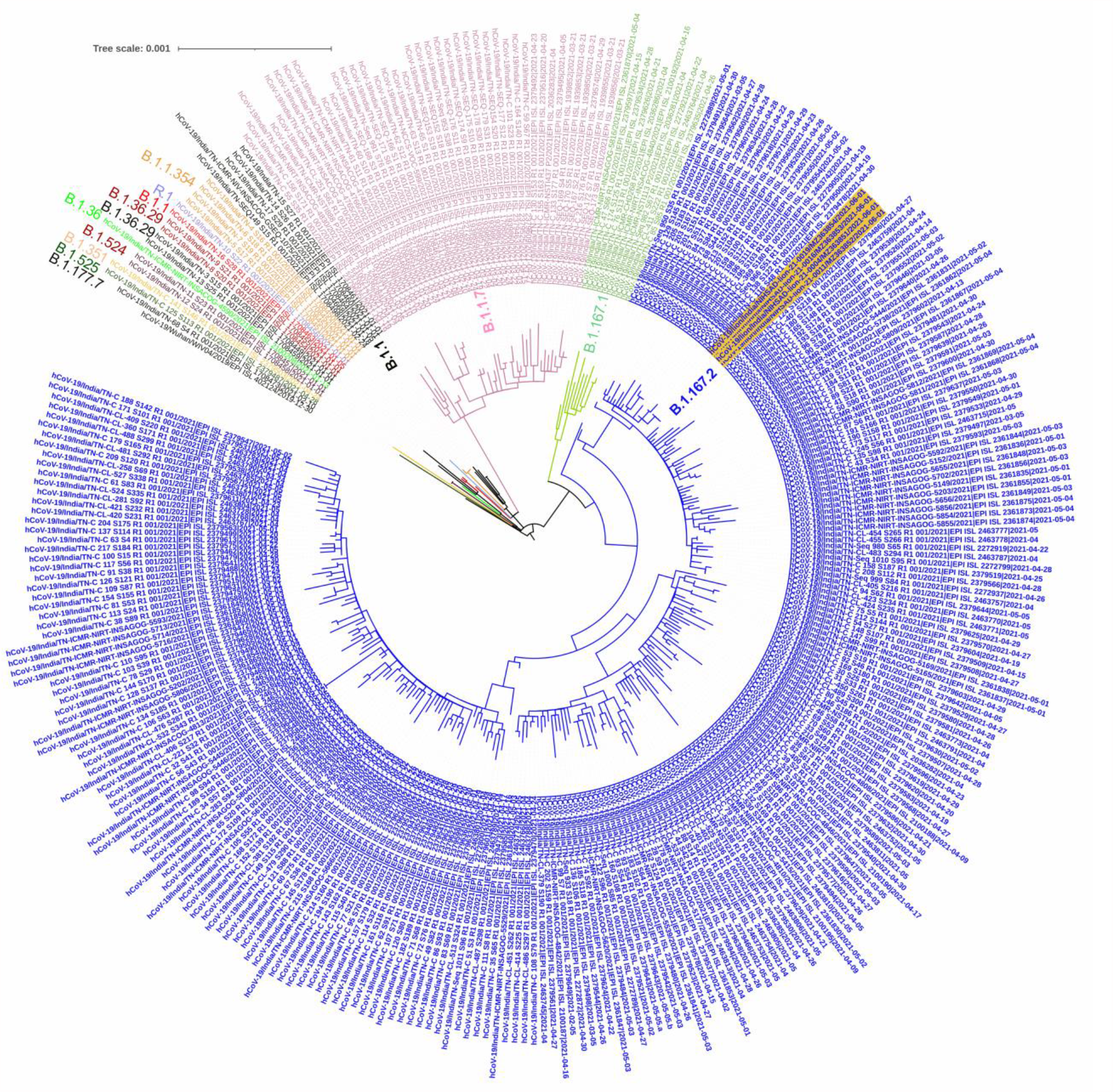
The phylogenetic analysis of Asiatic lions SARS-COV-2 sequences with all the available SARS-COV-2 sequences from Tamil Nadu state of India. The PANGO lineages were differently colored.

**Table S1.** Details of samples collected from 11 Asiatic lions in Arignar Anna Zoological Park, Chennai, Tamil Nadu, India.

| S. No | Identity of Lions | Age (years) | Type of sample | RT-qPCR Results (C<sub>t</sub> Value) |  |  | Results |
| --- | --- | --- | --- | --- | --- | --- | --- |
|  |  |  |  | E gene | RdRp | N |  |
| 1. | Lioness- Jeya | 3 | Nasal Swab | 20.16 | 19.65 | 19.97 | Positive |
|  |  |  | Rectal Swab | 19.23 | 18.27 | 19.75 | Positive |
| 2. | Lion- Padmnabhan | 12 | Nasal Swab | 18.90 | 18.10 | 18.50 | Positive |
|  |  |  | Rectal Swab | 27.41 | 26.75 | 27.79 | Positive |
| 3. | Lioness- Kavitha | 18 | Nasal Swab | 23.79 | 22.87 | 23.89 | Positive |
|  |  |  | Rectal Swab | 32.19 | 31.44 | 31.90 | Positive |
|  |  |  | Faecal sample | 27.56 | 27.20 | 27.72 | Positive |
| 4. | Lion- Shankar | 18 | Nasal Swab | 19.26 | 18.66 | 18.82 | Positive |
|  |  |  | Rectal Swab | 30.22 | 29.41 | 29.45 | Positive |
| 5. | Lioness- Neela | 9 | Nasal Swab | 22.71 | 23.22 | 22.57 | Positive |
|  |  |  | Rectal Swab | 25.09 | 24.74 | 24.46 | Positive |
|  |  |  | Faecal sample | - | - | - | Negative |
| 6. | Lioness- Niranjana | 2 | Nasal Swab | 19.68 | 19.04 | 19.17 | Positive |
|  |  |  | Rectal Swab | 38.61 | 38.14 | 33.69 | Negative |
|  |  |  | Faecal sample | - | - | - | Negative |
| 7. | Lion- Pradeep | 2 | Nasal Swab | 17.59 | 16.85 | 17.35 | Positive |
|  |  |  | Rectal Swab | 29.05 | 28.60 | 29.02 | Positive |
|  |  |  | Faecal sample | - | - | - | Negative |
| 8. | Lion- Vishnu | 4 | Faecal sample | 24.34 | 23.80 | 24.72 | Positive |
|  |  |  | Nasal swab | 26.60 | 25.43 | 26.74 | Positive |
| 9. | Lioness- Jeya@Bhuvana | 12 | Faecal sample | 28.11 | 27.39 | 28.49 | Positive |
|  |  |  | Nasal swab | 27.06 | 26.11 | 27.36 | Positive |
| 10. | Lion- Veera | 10 | Faecal sample | - | - | - | Negative |
| 11. | Lion- Shiva | 12 | Nasal swab | - | - | - | Negative |
|  |  |  | Throat swab | - | - | - | Negative |
|  |  |  | Trachea sample | - | - | - | Negative |
|  |  |  | Lymph node sample | - | - | - | Negative |
|  |  |  | Lungs sample | - | - | - | Negative |

**Table S2.**
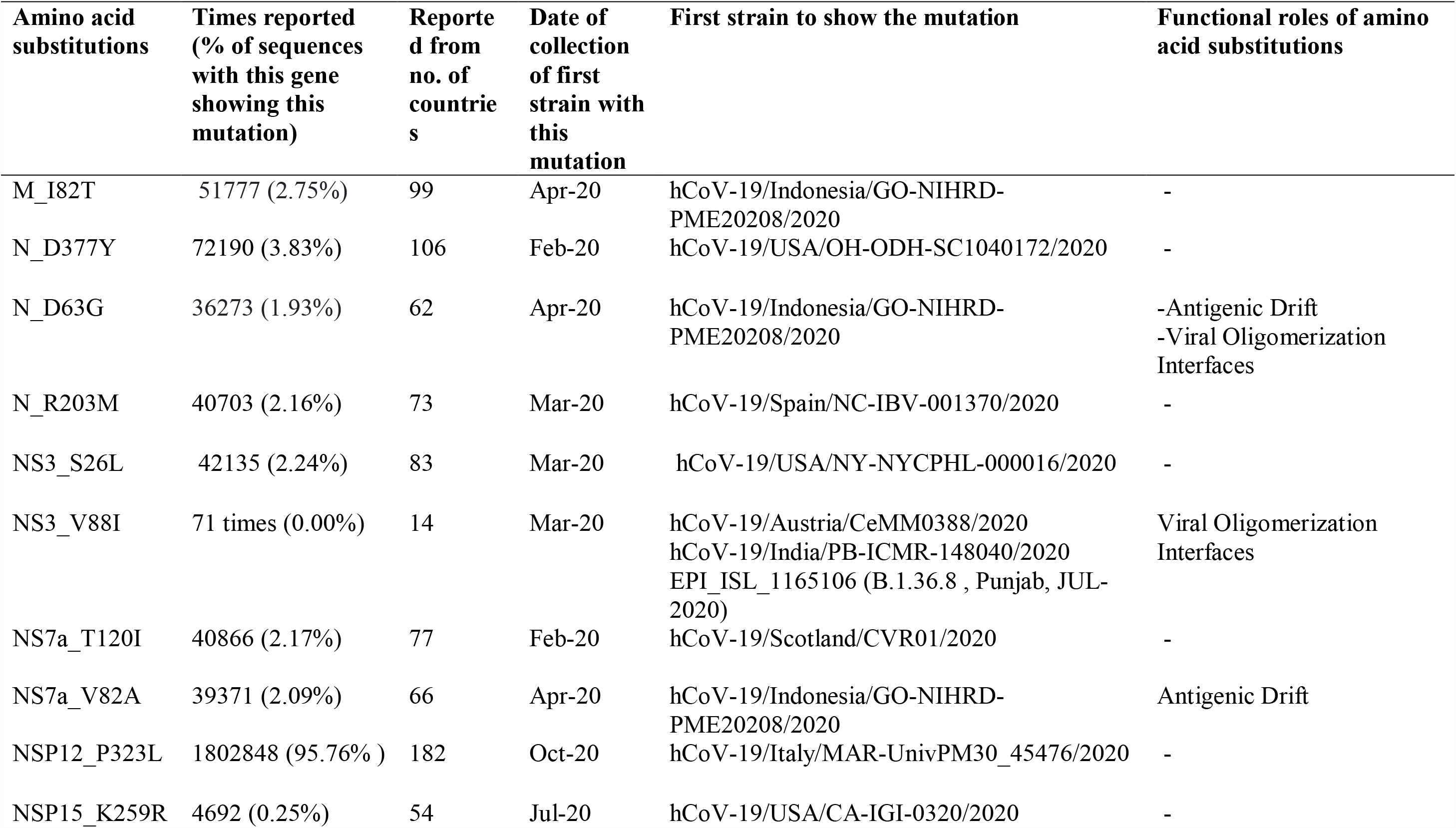
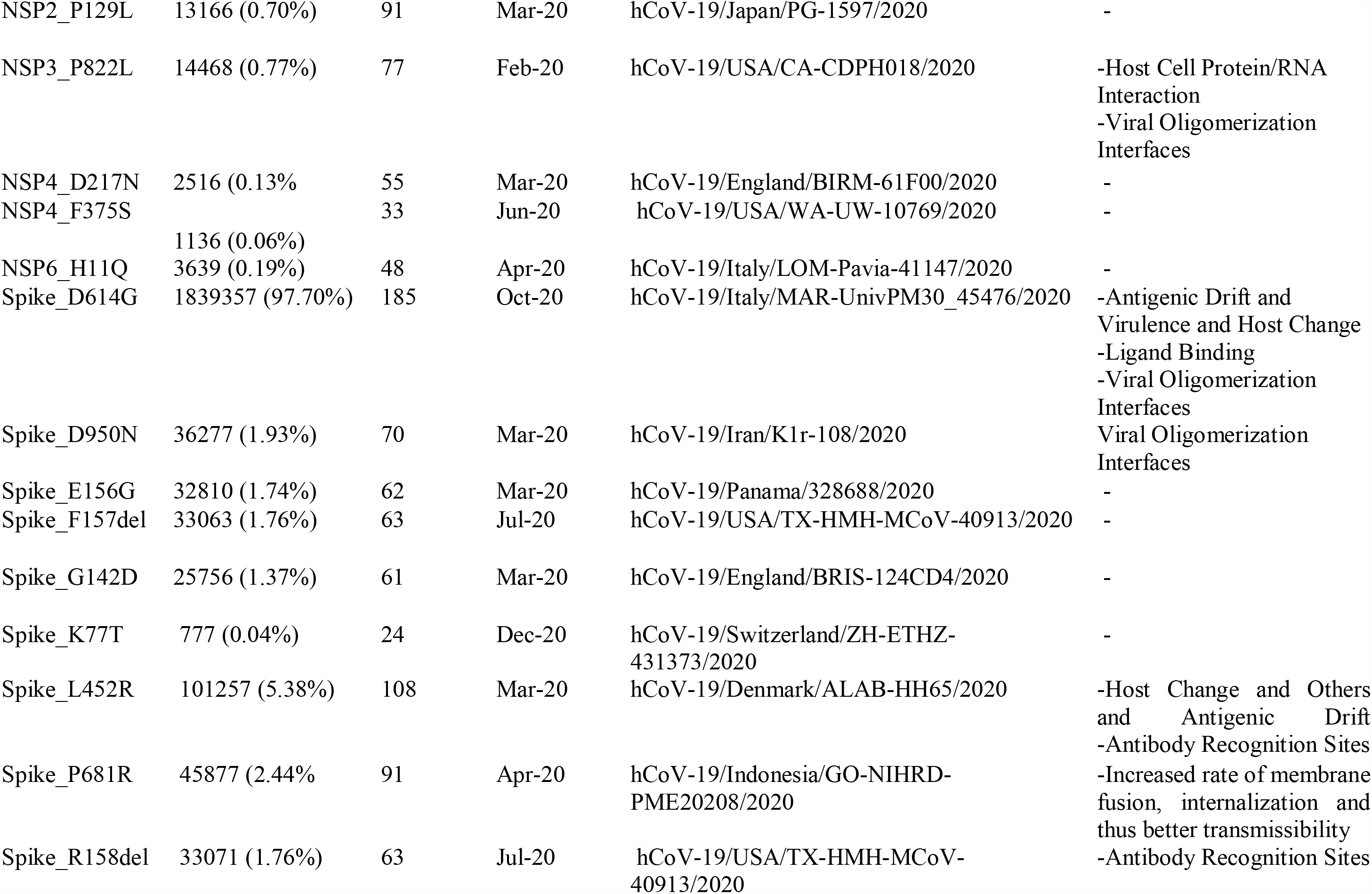
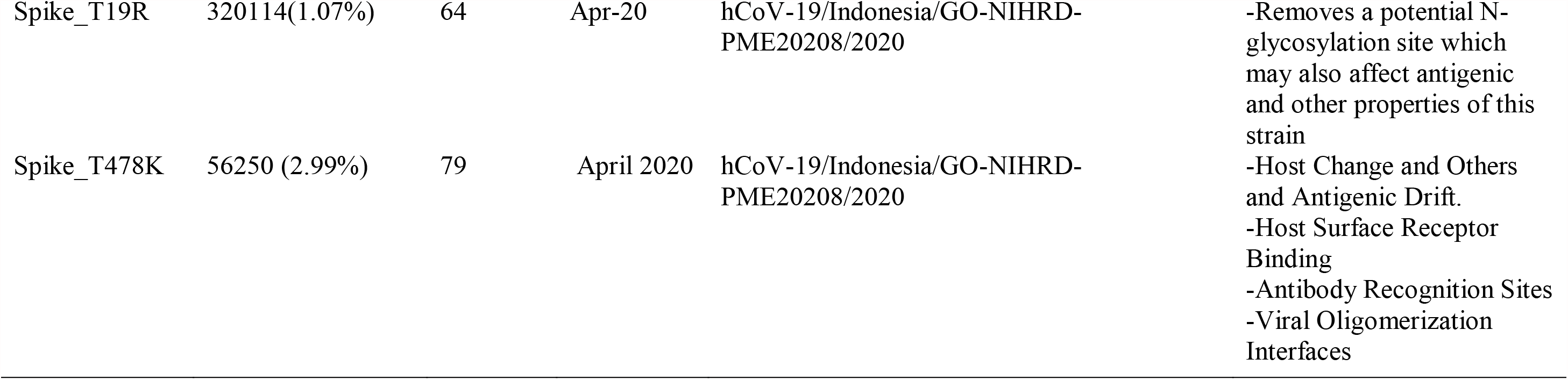
Amino acid substitutions identified in different proteins encoded by the SARS-CoV-2 of lions in comparison to Wuhan-Hu-1 reference sequence (GISAID accession no. EPI_ISL_402124), along with their functional roles.

## References

1. Bauer, H., Packer, C., Funston, P.F., Henschel, P. & Nowell, K. 2016. Panthera leo (errata version published in 2017). The IUCN Red List of Threatened Species 2016: e.T15951A115130419. https://dx.doi.org/10.2305/IUCN.UK.2016-3.RLTS.T15951A107265605.en. Downloaded on 16 June 2021.

2. Edgar RC. Search and clustering orders of magnitude faster than BLAST. Bioinformatics. 2010 Oct 1;26(19):2460–1. doi: 10.1093/bioinformatics/btq461.

3. Elbe S, Buckland-Merrett G. Data, disease and diplomacy: GISAID’s innovative contribution to global health. Glob Chall. 2017 Jan 10;1(1):33–46. doi: 10.1002/gch2.1018.

4. Frisk AL, König M, Moritz A, Baumgärtner W. Detection of canine distemper virus nucleoprotein RNA by reverse transcription-PCR using serum, whole blood, and cerebrospinal fluid from dogs with distemper. J Clin Microbiol. 1999 Nov;37(11):3634–43. doi: 10.1128/JCM.37.11.3634-3643.1999.

5. Katoh K, Standley DM. MAFFT multiple sequence alignment software version 7: improvements in performance and usability. Mol Biol Evol. 2013 Apr;30(4):772–80. doi: 10.1093/molbev/mst010.

6. Kim YI, Kim SG, Kim SM, Kim EH, Park SJ, Yu KM, Chang JH, Kim EJ, Lee S, Casel MAB, Um J, Song MS, Jeong HW, Lai VD, Kim Y, Chin BS, Park JS, Chung KH, Foo SS, Poo H, Mo IP, Lee OJ, Webby RJ, Jung JU, Choi YK. Infection and Rapid Transmission of SARS-CoV-2 in Ferrets. Cell Host Microbe. 2020 May 13;27(5):704-709.e2. doi: 10.1016/j.chom.2020.03.023.

7. Lam TT, Jia N, Zhang YW, Shum MH, Jiang JF, Zhu HC, Tong YG, Shi YX, Ni XB, Liao YS, Li WJ, Jiang BG, Wei W, Yuan TT, Zheng K, Cui XM, Li J, Pei GQ, Qiang X, Cheung WY, Li LF, Sun FF, Qin S, Huang JC, Leung GM, Holmes EC, Hu YL, Guan Y, Cao WC. Identifying SARS-CoV-2-related coronaviruses in Malayan pangolins. Nature. 2020 Jul;583(7815):282–285. doi: 10.1038/s41586-020-2169-0.

8. Li H. Minimap2: pairwise alignment for nucleotide sequences. Bioinformatics. 2018 Sep 15;34(18):3094–3100. doi: 10.1093/bioinformatics/bty191.

9. Liu P, Jiang JZ, Wan XF, Hua Y, Li L, Zhou J, Wang X, Hou F, Chen J, Zou J, Chen J. Are pangolins the intermediate host of the 2019 novel coronavirus (SARS-CoV-2)? PLoS Pathog. 2020 May 14;16(5):e1008421. doi: 10.1371/journal.ppat.1008421.

10. Loman NJ, Quick J, Simpson JT. A complete bacterial genome assembled de novo using only nanopore sequencing data. Nat Methods. 2015 Aug;12(8):733–5. doi: 10.1038/nmeth.3444.

11. McAloose D, Laverack M, Wang L, Killian ML, Caserta LC, Yuan F, Mitchell PK, Queen K, Mauldin MR, Cronk BD, Bartlett SL, Sykes JM, Zec S, Stokol T, Ingerman K, Delaney MA, Fredrickson R, Ivančić M, Jenkins-Moore M, Mozingo K, Franzen K, Bergeson NH, Goodman L, Wang H, Fang Y, Olmstead C, McCann C, Thomas P, Goodrich E, Elvinger F, Smith DC, Tong S, Slavinski S, Calle PP, Terio K, Torchetti MK, Diel DG. From People to Panthera: Natural SARS-CoV-2 Infection in Tigers and Lions at the Bronx Zoo. mBio. 2020 Oct 13;11(5):e02220–20. doi: 10.1128/mBio.02220-20.

12. OIE, 2021. SARS-COV-2 IN ANIMALS – SITUATION REPORT 1. Available at: https://www.oie.int/app/uploads/2021/06/sars-cov-2-situation-report-1.pdf, accessed on 16th June, 2021

13. Oude Munnink BB, Sikkema RS, Nieuwenhuijse DF, Molenaar RJ, Munger E, Molenkamp R, van der Spek A, Tolsma P, Rietveld A, Brouwer M, Bouwmeester-Vincken N, Harders F, Hakze-van der Honing R, Wegdam-Blans MCA, Bouwstra RJ, GeurtsvanKessel C, van der Eijk AA, Velkers FC, Smit LAM, Stegeman A, van der Poel WHM, Koopmans MPG. Transmission of SARS-CoV-2 on mink farms between humans and mink and back to humans. Science. 2021 Jan 8;371(6525):172–177. doi: 10.1126/science.abe5901.

14. Rambaut A, Holmes EC, O’Toole Á, Hill V, McCrone JT, Ruis C, du Plessis L, Pybus OG. A dynamic nomenclature proposal for SARS-CoV-2 lineages to assist genomic epidemiology. Nat Microbiol. 2020 Nov;5(11):1403–1407. doi: 10.1038/s41564-020-0770-5.

15. Shi J, Wen Z, Zhong G, Yang H, Wang C, Huang B, Liu R, He X, Shuai L, Sun Z, Zhao Y, Liu P, Liang L, Cui P, Wang J, Zhang X, Guan Y, Tan W, Wu G, Chen H, Bu Z. Susceptibility of ferrets, cats, dogs, and other domesticated animals to SARS-coronavirus 2. Science. 2020 May 29;368(6494):1016–1020. doi: 10.1126/science.abb7015.

16. Stamatakis A. RAxML version 8: a tool for phylogenetic analysis and post-analysis of large phylogenies. Bioinformatics. 2014 May 1;30(9):1312–3. doi: 10.1093/bioinformatics/btu033.

17. World Health Organization (2021). Available at: https://covid19.who.int/ (accessed on 17 June, 2012)

18. Zhang T, Wu Q, Zhang Z. Probable Pangolin Origin of SARS-CoV-2 Associated with the COVID-19 Outbreak. Curr Biol. 2020 Apr 6;30(7):1346-1351.e2. doi: 10.1016/j.cub.2020.03.022.

19. Zhou P, Yang XL, Wang XG, Hu B, Zhang L, Zhang W, Si HR, Zhu Y, Li B, Huang CL, Chen HD, Chen J, Luo Y, Guo H, Jiang RD, Liu MQ, Chen Y, Shen XR, Wang X, Zheng XS, Zhao K, Chen QJ, Deng F, Liu LL, Yan B, Zhan FX, Wang YY, Xiao GF, Shi ZL. A pneumonia outbreak associated with a new coronavirus of probable bat origin. Nature. 2020 Mar;579(7798):270–273. doi: 10.1038/s41586-020-2012-7.

